# Atto 465 derivative is a nuclear stain with unique excitation and emission spectra for multiplex immunofluorescence histochemistry

**DOI:** 10.1101/2021.05.14.444234

**Authors:** Joshua T. Dodge, Ana C. Costa-da-Silva, Christopher T. Hogden, Eva Mezey, Jacqueline W. Mays

## Abstract

The architecture of a biologic response is inextricably linked with the tissue architecture of the target site. Multiplex immunofluorescence (mIF) is an effective technique for the maximal visualization of multiple target proteins in situ. This powerful tool is limited by the spectral overlap separation of the currently available synthetic fluorescent dyes. The fluorescence excitation wavelengths ranging between 405nm and 488nm are rarely used in mIF imaging and serve as a logical additional slot for a fluorescent probe. In the present study, we demonstrate that Atto 465-pentafluoroaniline (Atto 465-p), a derivative of the fluorescent dye Atto 465, can serve as a nuclear stain in the violet-blue region of the visible spectrum. This opens the 405nm channel, traditionally used for nuclear stain, for detection of another experimental target. This increases the flexibility of the mIF panel and, with appropriate staining and microscopy, enables the quantitative analysis of at least six targets in one tissue section.

## Introduction

Recent advances in multiplex immunofluorescence (mIF) technology allow the simultaneous detection of multiple target proteins on a single tissue sample. This provides an accurate characterization of different cellular types and states in the context of their spatial relationship within the local environment. Diagnosis, treatment, and a better understanding, of complex diseases, such as cancer (1-3) and neurogenerative diseases (4, 5) is made possible via mIF. This sensitive technique can be affected by many factors including cross-reactivity of secondary antibodies and the spectral overlap of fluorophores. Several methods were recently developed to circumvent these limitations and achieve accurate mIF staining (6, 7). However, some require expensive, specialized kits and equipment (7, 8).

The tyramide signal amplification (TSA) system is based on a horseradish peroxidase (HRP) catalyzed deposition of fluorophore-labelled tyramide on and near a target protein or nucleic acid (9, 10). This straightforward technique is highly sensitive and specific, enables the precise detection of low-abundance proteins and the use of same-species antibodies (11, 12) and is cost-effective. Although a variety of target species and fluorophores are commercially available (12), it is rare for researchers to make use of the functional excitation wavelengths between 405 and 488 nm.

Atto 465 NHS ester, hereafter referred to as Atto 465, is structurally similar to proflavine (13). During the course of work to use Atto 465 to label antibodies for mIF panels, it became apparent that the dye was acting instead as an effective nuclear stain. In this study, we demonstrate that Atto 465 stains nuclei and is cleanly imaged at 465nm. It’s derivative, Atto 465-p, has higher photostability than Atto 465 NHS ester and can be easily separated from both 405nm and 488nm wavelengths using a confocal microscope with a tunable excitation source. We also demonstrate its utility by using Atto 465-p as a nuclear probe in a 6-plex IF assay.

## Materials and Methods

### Animals

BALB/cJ female mice (8 – 12 weeks old) were purchased from The Jackson Laboratory (Bar Harbor, ME). All animal procedures were approved by the Institutional Animal Care and Use Committee of NIDCR under an approved animal protocol and were performed according to the guidelines of the institute.

### Human tissues

Human minor salivary gland (MSG) and oral mucosa (OM) were obtained from healthy volunteers enrolled on registered, IRB-approved protocol NCT03602599 in accordance with the Declaration of Helsinki. Formalin-fixed paraffin-embedded (FFPE) tonsils, kidney, and skin blocks were obtained from a commercial tissue bank. All tissues were fixed in formalin, embedded in paraffin, and serially cut into 5 µm thick sections onto charged slides. Human karyotype slides were donated from the National Human Genome Research Institute Cytogenetic and Microscopy Core.

### Atto 465 NHS ester – crosslinking reaction

Atto 465 NHS ester (Sigma Aldrich, Saint Louis, MO, Pub-Chem Substance ID 329757879) was resuspended in DMSO, aliquoted and lyophilized. After resuspension in 1% triethylamine DMSO, 2,3,4,5,6-Pentafluoroaniline, Sigma-Aldrich) was added to the lyophilized dye in 10% molar excess. Each reaction was brought to a 16mM concentration of Atto 465-NHS ester and stirred at room temperature (RT) overnight on a vibrating plate protected from light. Dye was diluted to a final volume of 50uL with DMSO, aliquoted and stored at -20oC.

### Steady-state absorption and fluorescence measurements

Absorption measurements were performed on a Nanophotometer N60 (Implen, Westlake Village, CA). Fluorescence emission spectra were measured on a SpectraMax i3 (Molecular Devices, San Jose, CA). The dye was diluted to 160 uM, and a blank with equal moles of reactants (minus the dye) was used. Samples were excited at 430 nm and the fluorescent intensity was measured from 455 – 650 nm. All data were normalized to the highest fluorescent or absorption value.

### Photobleaching, photoconverting, and imaging

FFPE human tonsils were deparaffinized in xylene substitute (Sigma-Aldrich, St. Louis, MO), rehydrated in a graded ethanol series and stained for 10 minutes with Atto 465-carboxcylic acid (8 uM; Sigma-Aldrich, St. Louis, MO), Atto 465-p (8 μM), or YoPro-1 (0.125 μM; Invitrogen, Grand Island, NY). Tissue sections were imaged using a Nikon A1R confocal microscope Nikon A1R confocal microscope (fitted with a Pan Fluor 40x/1.30 oil objective) using the NIS Elements imaging software (Nikon Instruments Inc., Melville, NY). Photobleaching was achieved through repetitive imaging every 34 seconds for 2.66 hours using a 486nm laser set to 60% power. Laser energy varies between microscopes, so the microscope settings were converted into energy density to ensure results could be reproduced by other microscopes. The energy density was calculated by measuring the laser power in millijoules (mJ) at the back focal plane of the objective, at the given pixel dwell and resolution, with a compact power and energy meter (Thorlabs Newton, NJ), then multiplying by the pixel dwell time (μs/pixel), to generate the total millijoules per pixel of each scan, or area per pixel (μm2/pixel) to get the final energy density per scan. This was performed similar to Doyle et. al (14). The confocal line average was set to 8 in Galvano mode to ensure the laser was constantly delivering power to the tissue throughout the 2.66 hours time-lapse. All images were captured at 1024×1024 pixels with a 1.1-pixel dwell time (μs/pixel). To capture the 405nm channel, the 405nm laser was set to 2% power. Thus, any effect the 405nm laser had in reversing the 486nm photo-bleaching was minimal. Autofocus was used to prevent focus drift over time. For photoconversion experiments, the 486nm laser was set to 3% power and a time-lapse was run for 12 hours with 20-minute imaging intervals.

### Zinc and formalin fixed tissue staining

Submandibular gland, oral mucosa, and lymph node from BALB/cJ mice and humans were harvested and either fixed for 48 h at RT in 10% neutral buffered formalin (Sigma-Aldrich, St. Louis, MO) or zinc-fixation buffer (pH 6.5 - 7), prepared as described (15). Tissues were processed, embedded in paraffin and serially cut at 5 µm thick sections. For staining, sections were deparaffinized in xylene substitute (Sigma-Aldrich, St. Louis, MO), and rehydrated in a decreasing ethanol series. Tissues were then stained with Atto 465-p (4 μM), Hoechst 33342, (16.2 μM; Invitrogen, Eugene, OR), and/or ToPro-3-3 (1 μM; Thermo-Fisher Scientific, Waltham, MA) for 10 minutes at RT, protected from light, followed by washes in phosphate buffered saline (PBS; Corning, Manassas, VA).

### Frozen section IHC

Mice lymph nodes were dissected, embedded in Optimum Cutting Temperature compound (OCT; Sakura, Torrance, CA) and flash-frozen in cooled 2-methylbutane (Sigma-Aldrich, St. Louis, MO). Five µm-thick cryosections were cut onto charged slides and stored at -80°C until use. For staining, sections were air-dried, rehydrated in PBS and fixed at -20oC for 5min in 1:1 mixture of methanol and acetone. After washing with PBS, tissues were stained with Atto 465-p (4 μM), Hoechst 33342 (16.2 μM), and/or ToPro-3-3, (1 μM) for 10 minutes at RT, protected from light, followed by serial washes with PBS.

### Mouse peritoneal exudate cell staining

BALB/cJ mice were euthanized, and cells were collected by peritoneal lavage with 5 mL ice-cold DMEM medium (Gibco, Life Technologies, Grand Island, NY). Peritoneal exudate cells were centrifuged at 200x g and 4oC, for 10 minutes. Total cell numbers were determined using an automated cell counter (Countess II FL, Thermo Fisher Scientific). Cells were plated onto eight-well chamber slides (Nunc, Rochester, NY) after resuspension in DMEM supplemented with 50 IU/mL penicillin G, 50 µg/mL streptomycin and 10% fetal calf serum at a density of 1 x 106 cells/mL. Cells were kept in an incubator at 37°C and 5% CO2 for 3 h, washed with Hank’s Balanced Salt Solution and fixed in 4% paraformaldehyde (PFA) for 30 minutes at RT. After washing with PBS, cells were stained with Atto 465-p (4 μM, 2 μM, 1 μM, or 0.5 μM), Hoechst 33342 (16.2 μM), and/or ToPro-3-3, (1 μM) for 10 minutes at RT, protected from light, followed by washes with PBS.

### mIF staining

mIF was performed as described previously (10) using tyramide signal amplification (TSA) and repeated removal of sequentially applied antibodies (leaving the fluorochrome labelled water insoluble tyramide behind) before the next staining. Briefly, FFPE tonsil specimens were deparaffinized with xylene substitute (Sigma-Aldrich) and rehydrated with decreasing concentrations of ethanol (100%, 95% and 70% vol/vol; Sigma-Aldrich). Antigen retrieval was performed by microwaving slides in 10 mM citric acid buffer (pH 6), and endogenous peroxidase (HRP) activity was blocked using Bloxall (Vector Labs, Burlingame, CA). Primary antibodies, appropriate HRP-conjugated polymers (R&D Systems, Minneapolis, MN) and tyramide-conjugated fluorophores, provided in Supplementary Tabel 1, were serially added to tissue sections, followed by microwave treatment and blocking steps after each cycle. Nuclei were counterstained with Atto 465-p at 4 μM for 10 minutes at RT. Slides mounted with Fluoro-Gel (Electron Microscopy Sciences, Hatfield, PA).

### Image acquisition and analysis

For Atto 465p titration and colocalization studies, tissues or cells were imaged using a Nikon A1R confocal microscope (fitted with a Pan Fluor 40x/1.30 oil objective) using the NIS Elements imaging software (Nikon Instruments Inc., Melville, NY).

mIF staining was imaged using a Leica SP8 confocal microscope (fitted with a HC APO CS2 40x/1.30 oil objective) using the Leica Application Suite X (LAS X) software Leica, Wetzlar, Germany). Images acquired from both Nikon and Leica microscopes were processed using Fiji is Just ImageJ (Fiji). Image colocalization was quantified using Volocity 3.6 software (Perkin Elmer, Waltham, MA). Threshold for each image was determined automatically according to the Costes method (16). Mander’s overlap coefficient was then created for each image, comparing Hoechst 33342 staining to both Atto 465 and ToPro-3-3.

### Statistical analysis

Statistical analysis was performed using GraphPad Prism 8.4.1 software (GraphPad). All values are expressed as mean ± SEM. Data were compared using repeated measures one-way ANOVA. P-value < 0.05 is considered statistically significant.

## Results

### Synthesis and characterization of Atto 465-p

Atto 465 is structurally similar to proflavine (13), however it has a smaller Stokes shift and an emission spectrum further away from the 488 channel (Figure S1). This allows Atto 465 to be more readily separated from the 488 channel than is proflavine (Figure S1 B-D). In the present work, we tested Atto 465-p, a derivative of the fluorescent dye Atto 465, as a nuclear probe. A schematic diagram of the nucleophilic substitution reaction with Atto 465-NHS ester and 2,3,4,5,6-pentafluoroaniline is proposed in Figure 1A (17). Atto 465-p absorption (left side) and fluorescence spectra (right side) in PBS are depicted in Figure 1B. The absorbance spectrum gave maxima at 439 and 465 nm and the fluorescence emission extended from 465 to beyond 550 nm wavelengths.

**Figure 1:**
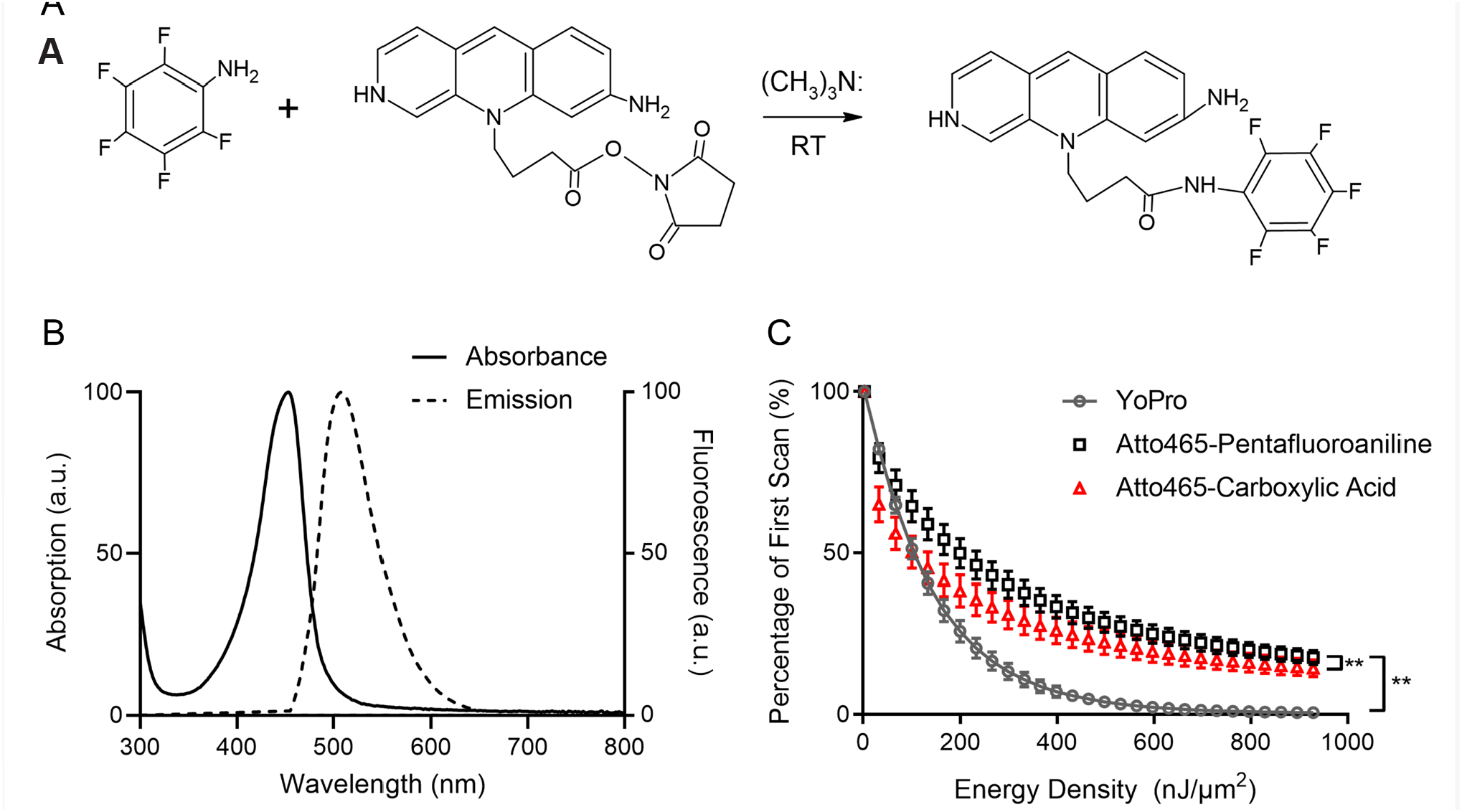
Characterization and photostability of Atto 465-p. A) Scheme illustrating presumed chemical structure and proposed nucleophilic substitution reaction between Atto 465-NHS ester and 2,3,4,5,6 pentafluoroaniline to synthetize Atto 465-p. B) Absorption (solid line) and fluorescence emission (dashed line) spectra of 8 µM Atto 465-p in PBS peaking at 450 nm and 508 nm, respectively. C) Photobleaching kinetics of YoPro-1, Atto 465-p and Atto 465 upon continuous irradiation by 486-nm laser. ** = p< 0.0001

revealing the marginal zone (MZ) and germinal center (GC) of a follicle. As expected, CD3, CD8 and cKit showed membranous staining, residing primarily extrafollicularly, whereas CD56 expression was membranous but present both in the extra and intrafollicular space. Moreover, IL-17 cytoplasmic staining was localized primarily outside the germinal center. As a proof of concept, each channel was evaluated individually showing that Atto 465-p can be fluorescently separated from the 405nm (IL-17 green) and 488nm (CD8 yellow) channels. Together, these data suggest that Atto 465-p can be added as a nuclear stain to a mIF protocol, thus facilitating the use of the 405 nm wavelength for another target.

## Discussion

In the present work, we demonstrate that Atto 465-p is a specific and stable fluorescent dye for nuclear staining with excellent performance. It can be used as an alternative to DAPI in mIF assays in mouse and human cells and tissues. These data show comparable nuclear staining between Atto 465-p and other common nuclear dyes Hoechst 33342 and ToPro-3.

Atto 465-p is a green fluorescent dye that absorbs light between 439 and 465 nm (Figure 1). It is easily excited with an argon laser in instruments emitting 488-nm light, which is a more common imaging component than is the UV laser. The Next, we evaluated Atto 465 free dye (carboxylic acid) and Atto 465-p for photobleaching behavior and compared them to a stable green nuclear dye, YoPro-1 (18). Under continuous excitation using 486 nm laser light, we observed faster reduction in fluorescence signal associated with YoPro-1 and Atto 465 free dye (carboxylic acid) when compared to Atto 465-p (Figure 1C), demonstrating that Atto 465-p has greater photostability. Further characterization revealed that Atto 465-p undergoes a photoconversion from green to a blue-emitting form upon exposure to a 486 nm light in a dosage dependent manner (Figure 2A and B). However, Atto 465-p regains fluorescence in a few hours, while maintaining the blue fluorescent form (Figure 2B).

**Figure 2:**
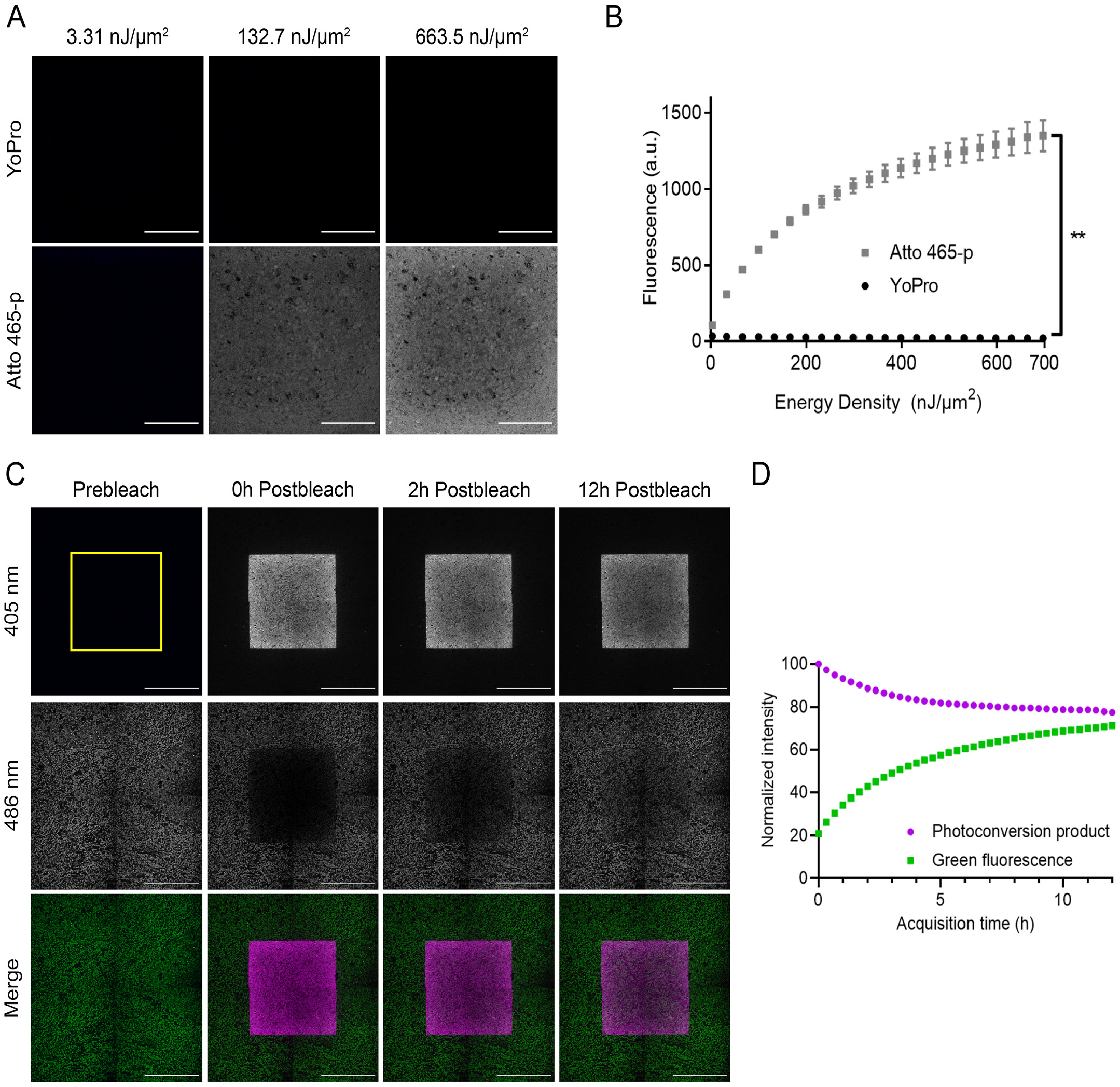
Characterization of Atto 465-p photoconversion. A) Representative images of human FFPE tonsil stained with YoPro-1 (top; 0.125μM) or Atto 465-p (bottom; 8μM) and subjected to photobleaching. Photoconversion product was detectable with illumination at 405nm. Scale bar=100μm. B) Plot illustrates the correlation between photoconverted fluorescence quantification and laser power density for both Atto 465-p (grey squares) and YoPro-1 (black circles). C) Representative time series images of human FFPE tonsil stained with Atto 465-p (8μM). The first column shows images taken prior to photobleaching (Prebleach); immediately after photobleaching (second column, 0h Postbleach), a 2-hour timepoint (third column, 2h Post-bleach) and 12-hour (fourth column, 12h Postbleach) timepoint using the same image settings as the pre-photobleached image (see Material and Methods). Scale bar=200μm. D) Graph shows the kinetic analysis of the average fluorescence intensities in the bleach ROI (yellow square in C) excited at 405 nm (magenta dots) and 486 nm (green squares) immediately after photobleaching (Time 0h). ** = p< 0.0001, paired t-test.

### Atto 465-p is a nuclei-specific fluorescent dye

We next evaluated the nuclear localization performance of Atto 465-p in multiple cell and tissue types. We determined an optimal working concentration of Atto 465-p from four different dilutions (0.5 µM to 4 µM) for fixed peritoneal exudate cells with 10-minute incubation. An efficient fluorescence signal was achieved at 4 µM as demonstrated in Figure 3A. We compared Atto 465-p staining to ToPro-3 and another common nuclear dye, Hoechst 33342. These dyes were selected because they differ in fluorescent spectrum, allowing them to be stained together on the same tissue. The resultant staining shows a similar pattern for Atto 465-p, Hoechst 33342 and ToPro-3 (Figure 3B). Single-color slides were stained to rule out fluorescent bleed through between channels (Figure 3B).

**Figure 3:**
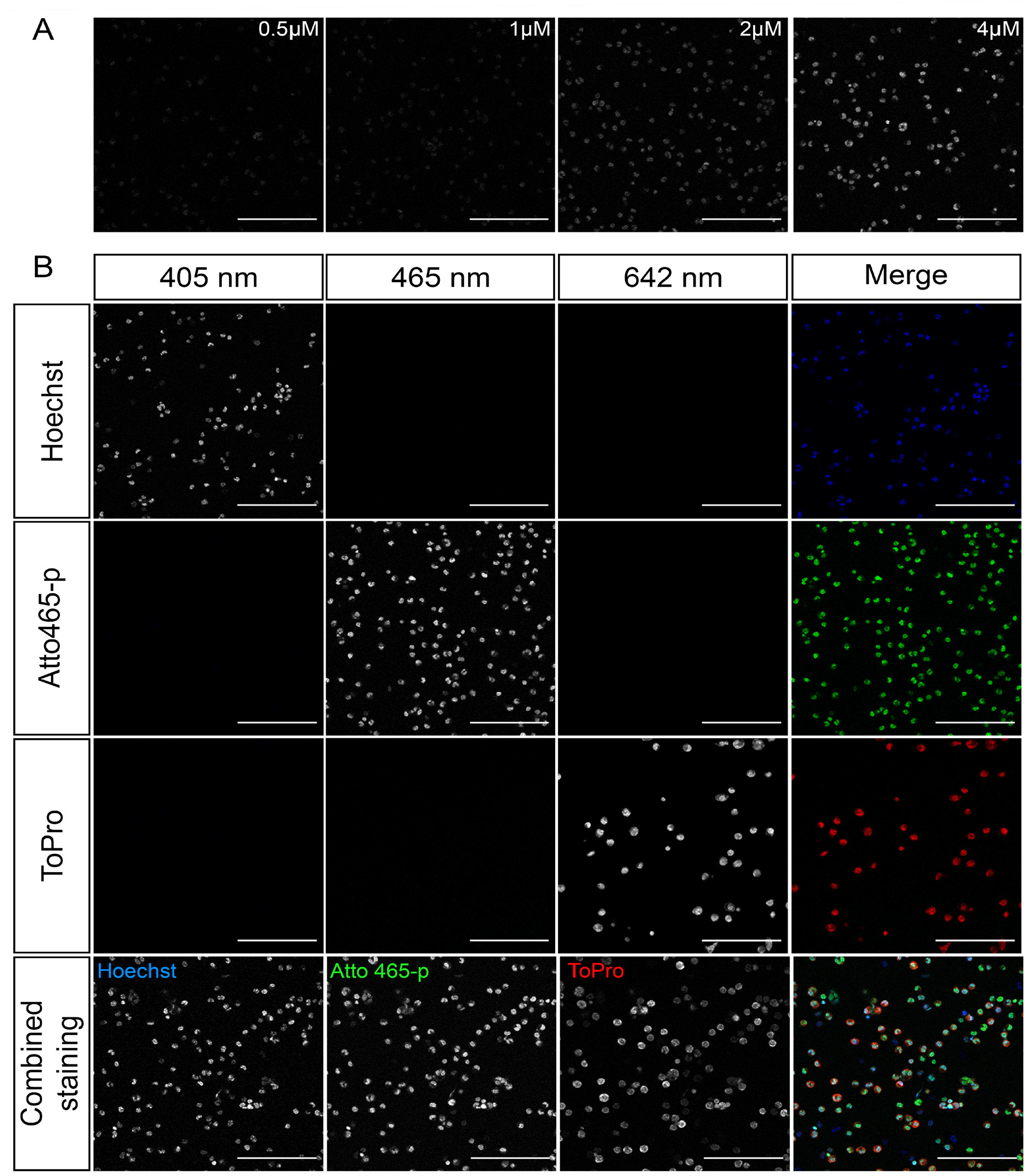
A) Serial dilution staining was done on 4%PFA fixed mouse peritoneal cells to determine optimal concentration for Atto 465-p. B) Mouse cells stained with either Hoechst 33342, Atto 465-p, ToPro-3, or a combination of all dyes. Scale bar = 50µm

To further confirm Atto 465-p affinity and specificity for nuclei, we conducted chromosomal staining using Hoechst 33342 as the positive control. Here, Atto 465-p demonstrated similar staining patterns with Hoechst 33342, suggesting that Atto 465-p stains chromatin (Fig. 4).

**Figure 4:**
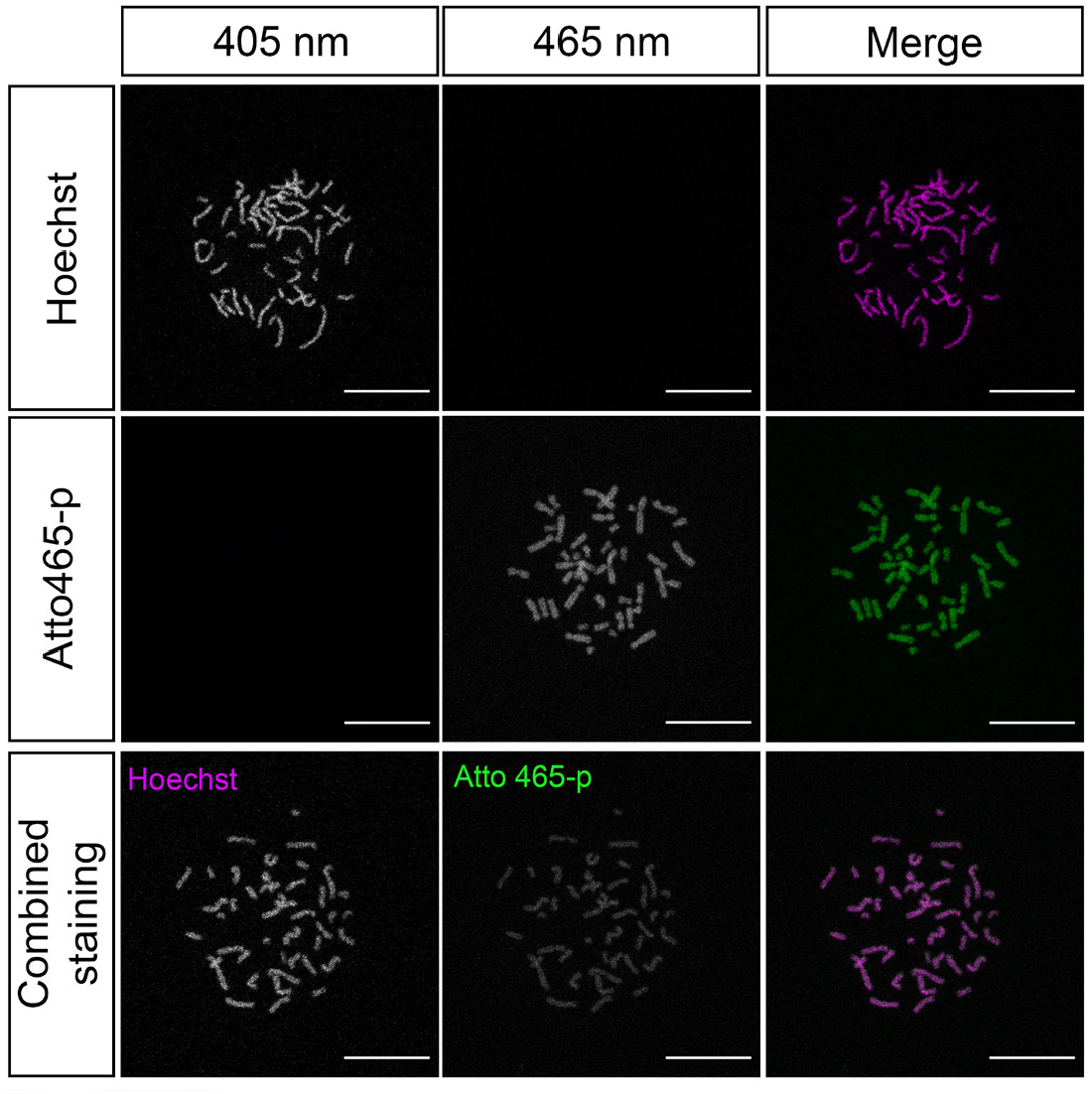
Immunofluorescence staining of human karyotype shows patterning of Hoechst 33342 and Atto 465-p with colocalization in the bottom row, suggesting that Atto 465-p and Hoechst 33342 label chromatin similarly. Scale bar = 25 µm

### Atto 465-p stains nuclei in various cell types and conditions, making available a novel fluorescent channel for mIF protocols

To further assess potential applications for Atto 465-p as a nuclear dye, several types of mouse (Figure 5A) and human (Figure S2) sections of tissues fixed with three disparate processing protocols were subsequently stained with Atto 465-p, Hoechst 33342 or ToPro-3,. Figures 5A and S2 show a nearly identical staining pattern of Atto 465-p, Hoechst 33342 and ToPro-3 in all tissues, independent of tissue fixation protocol. Quantitative colocalization analysis (Figure 5B and Supplementary Table 2) demonstrated higher degree of colocalization between Hoechst 33342 and Atto 465-p (R = 0.8043 +/-0.022) than Hoechst 33342 and ToPro-3 (R = 0.7242 +/-0.022).

**Figure 5:**
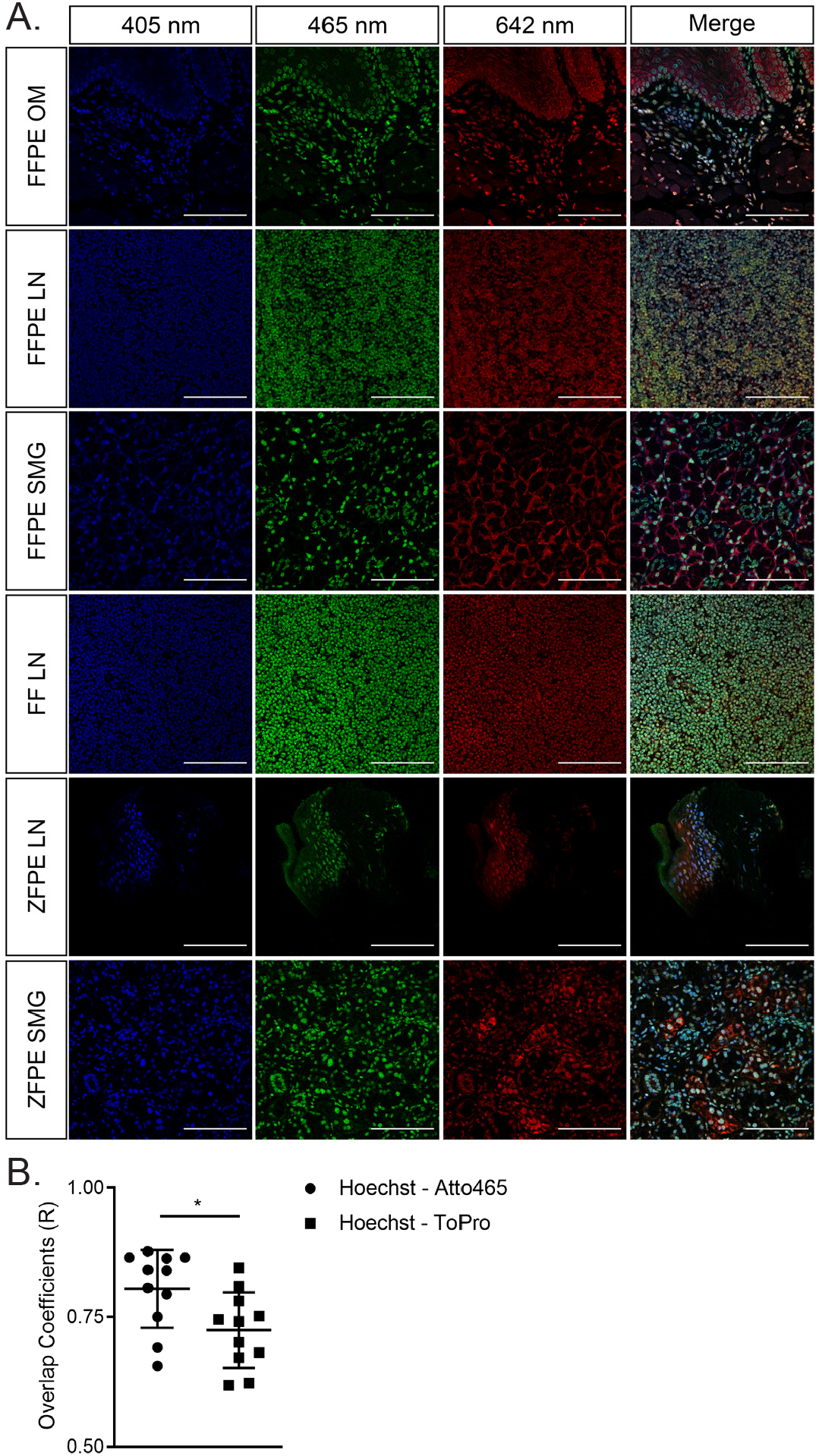
A) Mouse tissues following different fixation protocols to compare nuclear specificity of Atto 465-p to Hoechst 33342 and ToPro-3. OM = Oral mucosa; LN = lymph nodes; SMG = submandibular gland; ZFPE = zinc-fixed paraffin-embedded; FF – flash-frozen; Scale bar = 100µm. B) Quantification of the overlap coefficient demonstrated higher similarity between Hoechst 33342 and Atto 465-p than Hoechst 33342 and ToPro-3. Values represent mean +/-SEM. * = p < 0.001, paired t-test.

To test if Atto 465-p is suitable for mIF, we developed a 6-plex TSA panel using Atto 465-p as a nuclear probe. Human tonsil was stained for known surface markers, CD3 (T cells), CD8 (T cell subset), cKit (mast/innate lymphoid cells) and CD56 (NK cells) as well as an intracellular cytokine, IL-17, and visualized by confocal microscopy (Figure 6). Atto 465-p staining (shown in blue) highlights the border of each cellular nucleus, revealing the marginal zone (MZ) and germinal center (GC) of a follicle. As expected, CD3, CD8 and cKit showed membranous staining, residing primarily extrafollicularly, whereas CD56 expression was membranous but present both in the extra and intrafollicular space. Moreover, IL-17 cytoplasmic staining was localized primarily outside the germinal center. As a proof of concept, each channel was evaluated individually showing that Atto 465-p can be fluorescently separated from the 405nm (IL-17 green) and 488nm (CD8 yellow) channels. Together, these data suggest that Atto 465-p can be added as a nuclear stain to a mIF protocol, thus facilitating the use of the 405 nm wavelength for another target.

**Figure 6:**
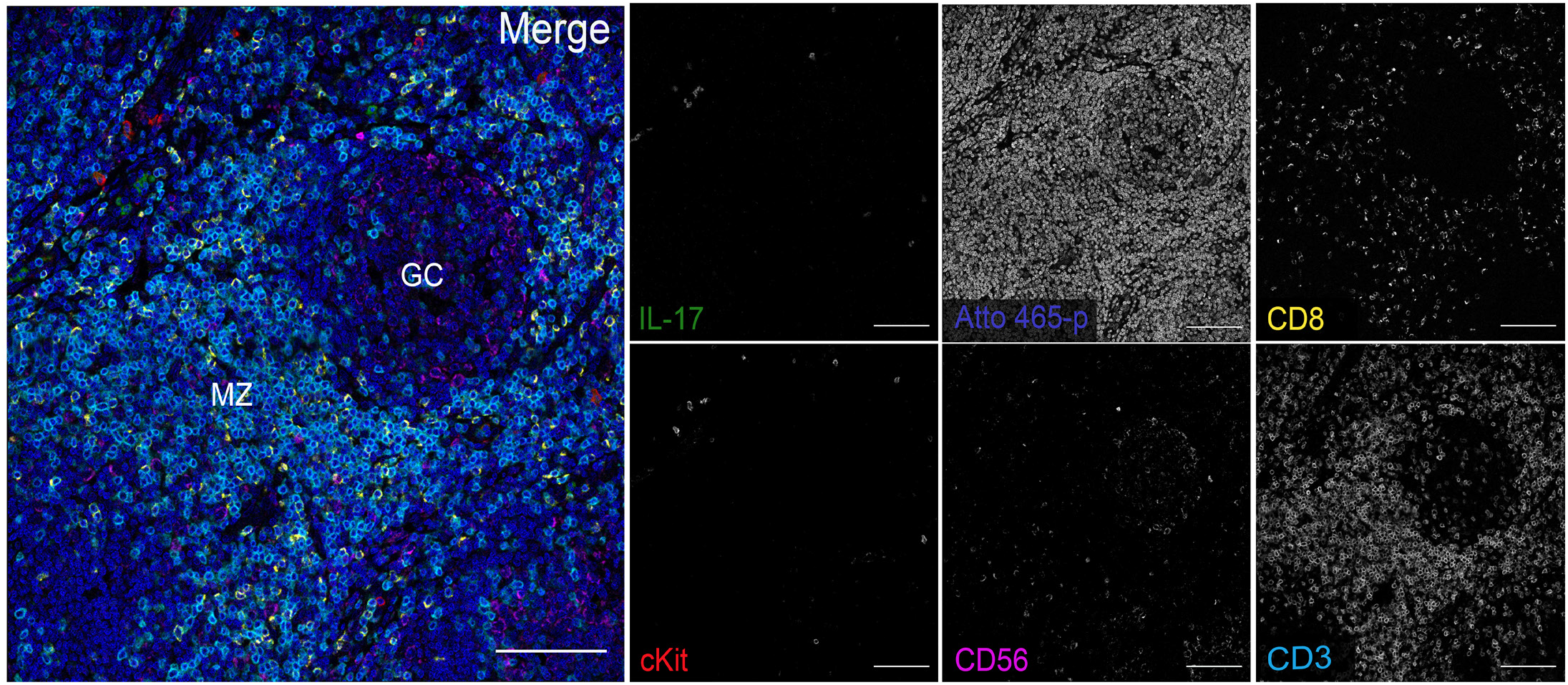
Atto 465-p as a nuclear dye. mIF microscopy of IL-17 (green), CD8 (yellow), cKit (red) CD56 (pink), and CD3 (cyan) in FFPE human tonsil. Atto 465-p was used as a nuclear probe and shows distinguishable staining from the 405 nm and 488 nm channels. GC = germinal center; MZ = marginal zone. Scale bar = 100 µm.

## Discussion

In the present work, we demonstrate that Atto 465-p is a specific and stable fluorescent dye for nuclear staining with excellent performance. It can be used as an alternative to DAPI in mIF assays in mouse and human cells and tissues. These data show comparable nuclear staining between Atto 465-p and other common nuclear dyes Hoechst 33342 and ToPro-3.

Atto 465-p is a green fluorescent dye that absorbs light between 439 and 465 nm (Figure 1). It is easily excited with an argon laser in instruments emitting 488-nm light, which is a more common imaging component than is the UV laser. The spectral properties of Atto 465-p favor its color separation from Alexa Fluor 405 and Alexa Fluor 488 (Figure 6), which provides a stable and specific alternative to leave the 405nm channel open for additional targets when spectral imaging or a white light laser are available.

Importantly, UV excitation can be more toxic to cells than visible light (19), which can hinder further applicability in live cell microscopy studies. Our results demonstrate that extended exposure to blue light (486nm) will prompt a photoconversion reaction whose product absorbs 405 nm light and emits blue fluorescence. Photoconversion has been shown in other common nuclear dyes like DAPI and Hoechst 33342 (20), although such event happened after high levels of energy were concentrated on the dye. A single image scan at normal settings is about 0.23 nJ/μm2, a much lower amount than what was used in these photobleaching experiments (Figure 2). While this photoconversion is a cautionary characteristic for IHC multiplexing, the same characteristic has been leveraged in experiments as a simple method to mark specific cells (21). It should be noted that the amount of energy used to bleach and photoconvert Atto 465p is much higher than typical confocal microscope settings for mIF. However, further investigation into the potential of Atto 465-p for live imaging is warranted. Leveraging of Atto 465-p as a photoinducible cell marker would be advantageous to the field, given that photoinducible markers, such as KikGR (22), PS-CFP (23), PA-GFP (24), PA-mRFP1 (25) and others, require ultraviolet light.

Taken together, Atto 465-p demonstrates both specificity and photostability and fulfils the requirements to be used as a nuclear dye for immunocytochemistry and immunohistochemistry applications. The use of Atto 465-p for nuclear staining increases the multiplex capacity in mIF assays by occupying an underutilized region in the fluorescent spectrum.

## Acknowledgements

This work was funded by the intramural program of the National Institute of Dental and Craniofacial Research, National Institutes of Health. Human karyotype slides were kindly prepared and provided by National Human Genome Research Institute Cytogenetic and Microscopy Core. This research was supported by the NIDCR Imaging Core: ZIC DE000750. The authors thank Dr. Andrew Doyle for his technical expertise and review of the manuscript.

## Supplementary Data

**Supplementary Table 1:** List of antibodies used for mIF

| Type | Antibody name and catalogue number | Dilution | Source | Cell Target |
| --- | --- | --- | --- | --- |
| Primary antibody | rat anti-CD3 | 1:2000 | BioRad, Hercules, CA | T cells |
|  | mouse anti-CD8 | 1:1000 | BioRad | T cell subset |
|  | rabbit anti-CD56 | 1:1000 | Abcam, Cambridge, MA | Natural killer cells |
|  | rabbit anti-cKit | 1:1000 | Abcam | Mast cell / Innate Lymphoid cell |
|  | goat anti-IL-17 | 1:1000 | R&D Systems, Minneapolis, MN | IL-17 producing cells |
| HRP-conjugated Tyramide antibodies | Alexa Fluor 405 | 1:500 | Thermo-Fisher Scientific, Eugene, OR | - |
|  | Alexa Fluor 488 | 1:200 |  | - |
|  | Alexa Fluor 555 | 1:100 |  | - |
|  | Alexa Fluor 594 | 1:100 |  | - |
|  | Atto 633 | 1:500 | Sigma-Aldrich, St. Louis, MO | - |

**Supplementary Table 2:** Overlap Coefficient – Figure 5

| Tissue | Hoechst 33342 to Atto 465-p Overlap Coefficient (R) | Hoechst 33342 to ToPro-3 Overlap Coefficient (R) |
| --- | --- | --- |
| Human FFPE Tonsil | 0.865 | 0.781 |
| Human FFPE MSG | 0.865 | 0.845 |
| Human FFPE Oral Mucosa | 0.691 | 0.622 |
| Human FFPE Skin | 0.750 | 0.701 |
| Human FFPE Kidney | 0.655 | 0.618 |
| Mouse FFPE Oral Mucosa | 0.794 | 0.752 |
| Mouse FFPE LN | 0.863 | 0.809 |
| Mouse FFPE SMG | 0.806 | 0.681 |
| Mouse Frozen Lymph Node | 0.877 | 0.745 |
| Mouse Zinc Lymph Node | 0.840 | 0.741 |
| Mouse Zinc SMG | 0.841 | 0.671 |

**Supplemental Figure 1:**
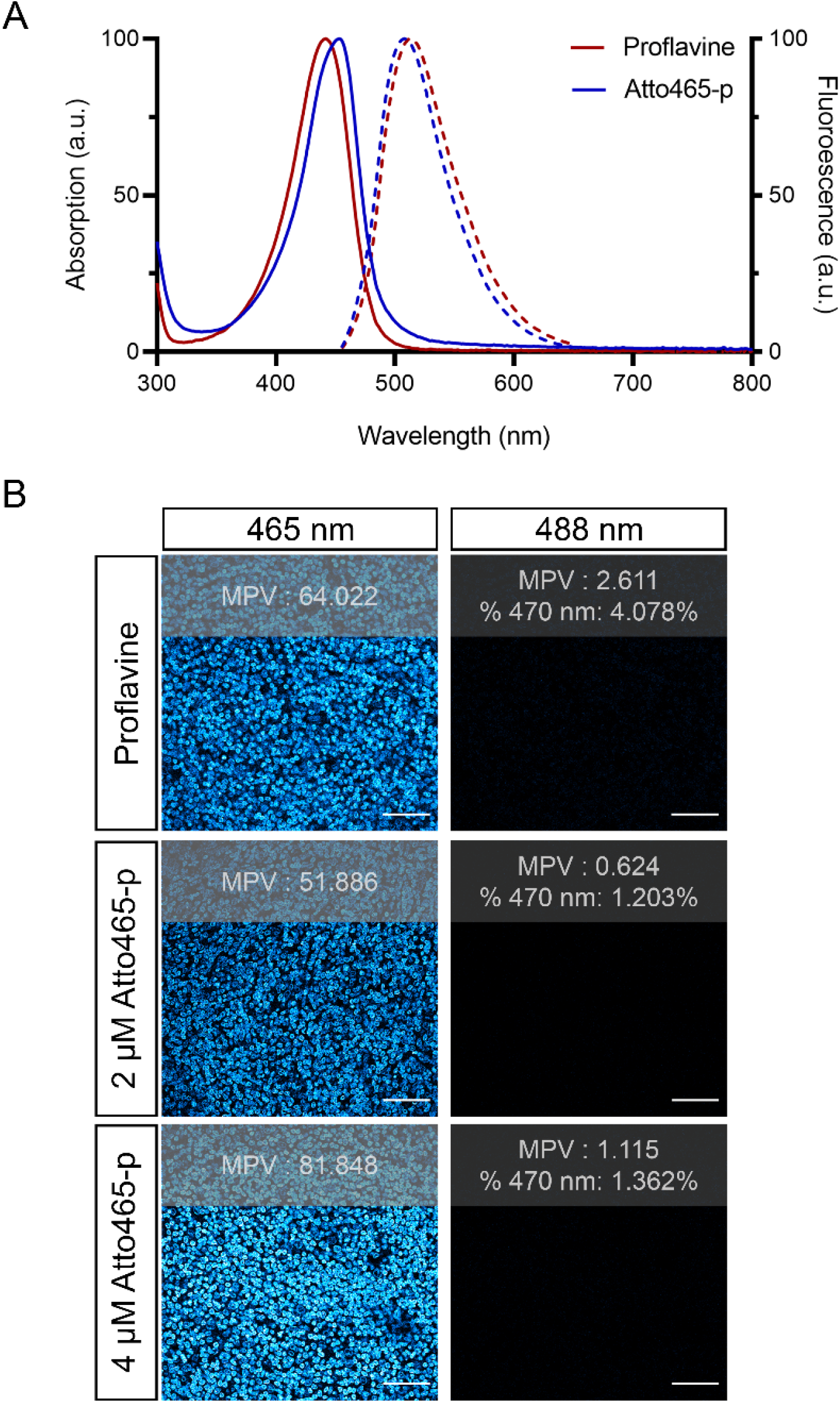
A) Absorption (solid line) and fluorescence emission (dashed line) spectra of 8 µM Proflavine (red line) and 8µM Atto 465-p (blue line). B) Human FFPE tonsil tissue was stained with 8 µM Proflavine, 2 µM or 4 µM of Atto 465-p fluorescence signal measured following excitation at 465 nm and 488 nm on a Leica SP8 using the same microscope settings. Images were converted to Cyan Hot lookup table (LUT) to better demonstrate staining intensities. Proflavine shows higher background and fluorescent bleed-through into 488 nm than Atto465-p. Scale bar = 25µm

**Supplemental Figure 2:**
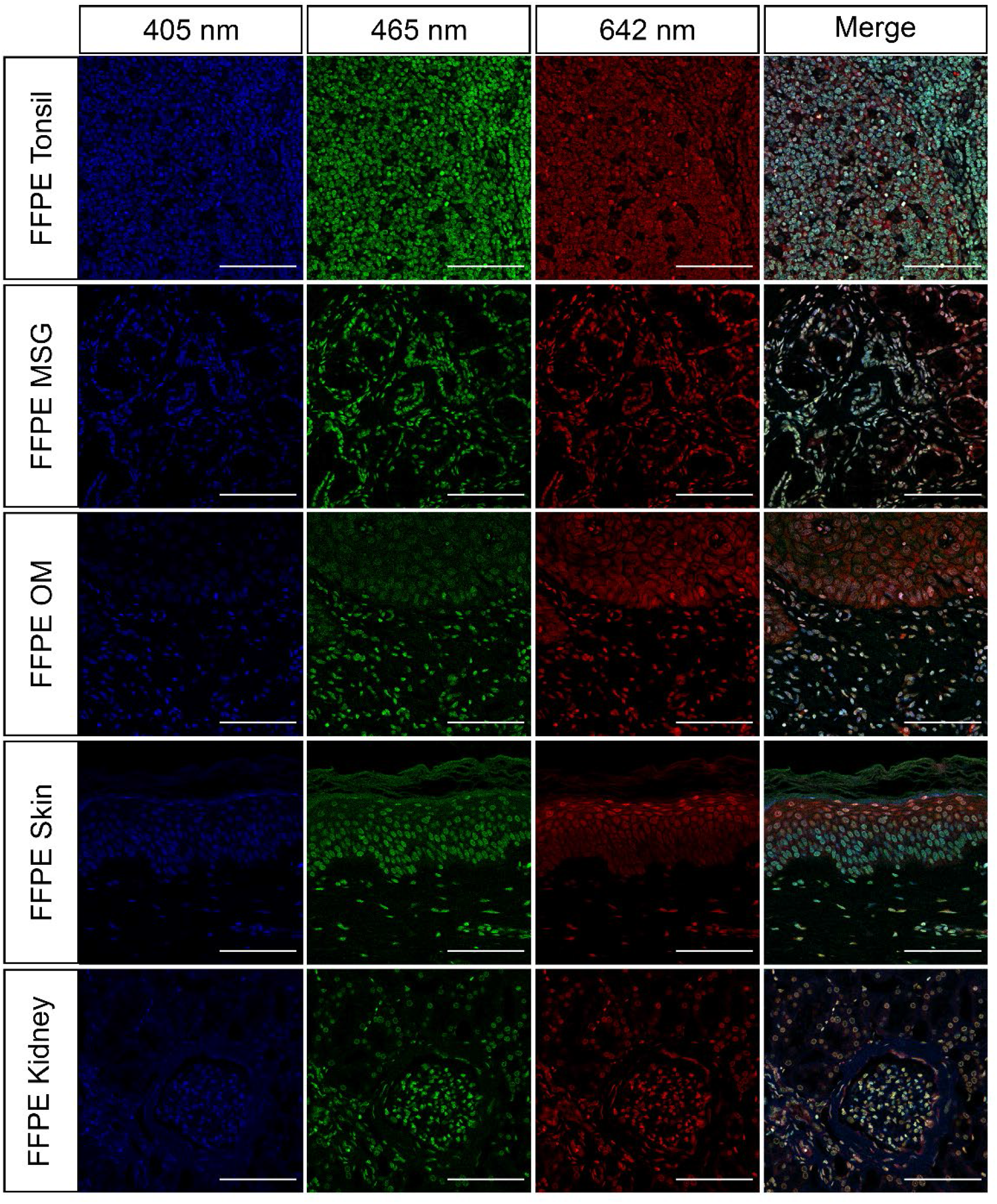
Human FFPE tonsil, MSG, OM, skin, and kidney were stained with Hoechst (16.2µM) and ToPro (1µM) and imaged on a Nikon confocal microscopy to demonstrate similar nuclear specificity of Atto 465-p (4 µM). Scale bar = 100µm.

